# Antibody responses to Omicron BA.4/BA.5 bivalent mRNA vaccine booster shot

**DOI:** 10.1101/2022.10.22.513349

**Authors:** Qian Wang, Anthony Bowen, Riccardo Valdez, Carmen Gherasim, Aubree Gordon, Lihong Liu, David D. Ho

## Abstract

The SARS-CoV-2 Omicron variant and its numerous sub-lineages have exhibited a striking ability to evade humoral immune responses induced by prior vaccination or infection. The Food and Drug Administration (FDA) has recently granted Emergency Use Authorizations (EUAs) to new bivalent formulations of the original Moderna and Pfizer mRNA SARS-CoV-2 vaccines that target both the ancestral strain as well as the Omicron BA.4/BA.5 variant. Despite their widespread use as a vaccine boost, little is known about the antibody responses induced in humans. Here, we collected sera from several clinical cohorts: individuals after three or four doses of the original monovalent mRNA vaccines, individuals receiving the new bivalent vaccines as a fourth dose, and individuals with BA.4/BA.5 breakthrough infection following mRNA vaccination. Using pseudovirus neutralization assays, these sera were tested for neutralization against an ancestral SARS-CoV-2 strain, several Omicron sub-lineages, and several related sarbecoviruses. At ~3-5 weeks post booster shot, individuals who received a fourth vaccine dose with a bivalent mRNA vaccine targeting BA.4/BA.5 had similar neutralizing antibody titers as those receiving a fourth monovalent mRNA vaccine against all SARS-CoV-2 variants tested, including BA.4/BA.5. Those who received a fourth monovalent vaccine dose had a slightly higher neutralizing antibody titers than those who received the bivalent vaccine against three related sarbecoviruses: SARS-CoV, GD-Pangolin, and WIV1. When given as a fourth dose, a bivalent mRNA vaccine targeting Omicron BA.4/BA.5 and an ancestral SARS-CoV-2 strain did not induce superior neutralizing antibody responses in humans, at the time period tested, compared to the original monovalent vaccine formulation.

## Main text

Continued evolution of Severe Acute Respiratory Coronavirus 2 (SARS-CoV-2), which causes Coronavirus Disease 2019 (COVID-19), has led to the emergence of the Omicron variant and numerous sub-lineages that evade neutralizing antibody responses induced by infection or vaccination^1^. In response to this concerning trend, the Food and Drug Administration granted Emergency Use Authorizations (EUAs) to bivalent formulations of mRNA vaccines produced by Pfizer and Moderna that target both the Omicron BA.4/BA.5 spike and an ancestral wild-type (WT) SARS-CoV-2 spike^2^. Published data on antibody responses to bivalent vaccines have been limited to animal studies and human studies utilizing a different bivalent mRNA vaccine targeting the Omicron BA.1 spike in addition to the WT spike^3,4^. Despite their widespread use, the impact of a booster shot with a new bivalent vaccine on SARS-CoV-2-neutralizing antibody responses in humans remains unknown.

Therefore, we collected a panel of sera from individuals who had received three doses of the original monovalent mRNA vaccines followed by one dose of a bivalent vaccine targeting BA.4/BA.5 (details in Supplementary Appendix). We compared virus neutralization by these sera to panels of sera from individuals who received either three or four monovalent mRNA vaccines as well as to sera from individuals with BA.4/BA.5 breakthrough infection following mRNA vaccination. Using pseudovirus neutralization assays, all sera were tested against an ancestral SARS-CoV-2 strain (D614G) and Omicron sub-lineages BA.1, BA.2, BA.4/BA.5, BA.4.6, BA.2.75, and BA.2.75.2. To further assess the breadth of antibody responses, we also tested sera for neutralization against several related sarbecoviruses: SARS-CoV, GD-pangolin, GX-pangolin, and WIV1.

Clinical details are summarized for all groups in **Table S1** and listed for each case in **Table S2**. Individuals who received four monovalent mRNA doses were older (mean age 55.3) than those who received a bivalent booster (mean age 36.4). Serum was collected from both cohorts at a similar time following the vaccine boost (mean 24.0 days in the monovalent group; mean 26.4 days in the bivalent group). All cohorts exhibited the highest serum virus-neutralization titers (ID_50_) against the ancestral D614G strain (**Figure 1A**). Geometric mean ID_50_ titers against SARS-CoV-2 variants were lowest for boosted sera and highest for BA.4/BA.5 breakthrough sera. There was no significant difference in neutralization of any SARS-CoV-2 variant tested between individuals who received a fourth monovalent vaccine and those who received a fourth dose of a bivalent vaccine (**Figure 1B**). ID_50_ titers against three related sarbecoviruses (SARS-CoV, GD-Pangolin, and WIV1) were slightly but significantly higher in those who received a fourth monovalent vaccine dose compared to those who received a bivalent vaccine.

**Figure 1.**
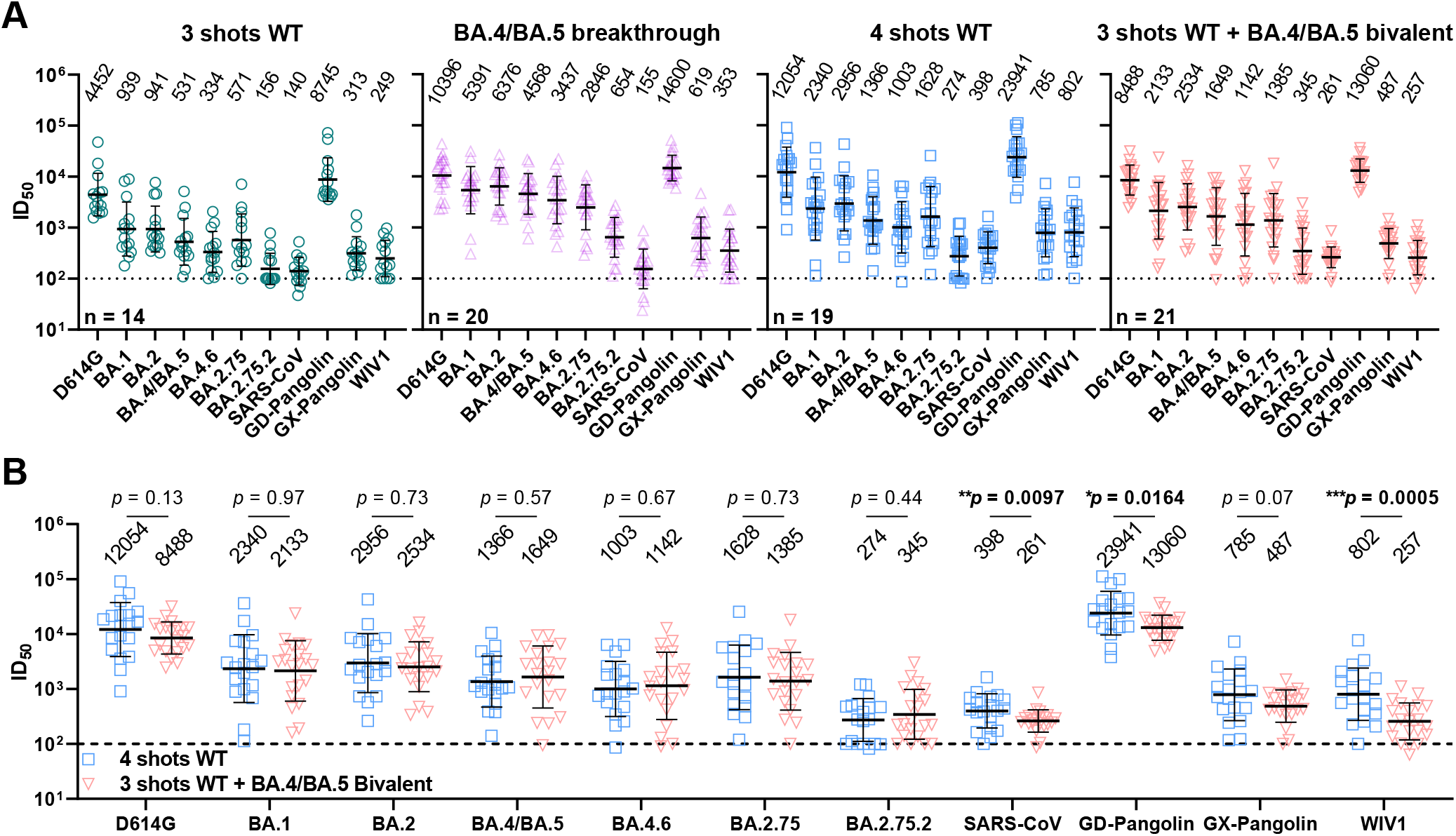
Neutralization profiles of serum samples against SARS-CoV-2 variants and other sarbecoviruses. **(A)** Neutralization ID_50_ titers of serum samples from “3 shots WT”, “BA.4/BA.5 breakthrough”, “4 shots WT”, and “3 shots WT + bivalent” cohorts. Values above the symbols denote the geometric mean ID_50_ titers, and values on the lower left indicate the sample size (n). The limit of detection is 100 (dotted line). Wild-type (WT) shots refer to monovalent mRNA vaccine doses. “3 shots WT,” sera from individuals vaccinated with three doses of the WT mRNA vaccine; “BA.4/BA.5 breakthrough,” sera from individuals with BA.4 or BA.5 infection following WT mRNA vaccination; “4 shots WT,” sera from individuals vaccinated with four doses of the WT mRNA vaccine; “3 shots WT + bivalent,” sera from individuals vaccinated with three doses of the WT mRNA vaccine and subsequently one dose of a BA.4/BA.5 bivalent mRNA vaccine. **(B)** Comparison of antibody responses induced by a fourth dose of the original WT mRNA vaccine versus a fourth dose of a BA.4/BA.5 bivalent mRNA vaccine. Comparisons were made by Mann-Whitney tests. \**p* < 0.05; \*\**p* < 0.01; \*\*\**p* < 0.001. Values above the symbols denote the geometric mean ID_50_ titers.

Boosting with a new bivalent mRNA vaccine targeting both BA.4/BA.5 and an ancestral SARS-CoV-2 strain did not elicit a discernibly superior virus-neutralizing antibody responses compared boosting with an original monovalent vaccine. These findings may be indicative of immunological imprinting^5^, although follow-up studies are needed to determine if the antibody responses will deviate in time, including the impact of a second bivalent booster.

