## Supplemental figures for "Antibody responses to Omicron BA.4/BA.5 bivalent mRNA vaccine booster shot"

### Supplementary Methods

*Clinical cohorts*

Sera analyzed in this study was categorized into several cohorts. Boosted samples consisted of sera from individuals who had received three doses of monovalent, referred to as wild-type (WT), mRNA vaccines (either Moderna mRNA-1273 or Pfizer BNT162b2). Sera was also collected from individuals after a fourth monovalent mRNA vaccine (referred to as “4 shots WT”). Bivalent vaccine sera were collected from individuals who had received three monovalent mRNA vaccine doses followed by one dose of the Pfizer or Moderna bivalent vaccine targeting BA.4/BA.5 in addition to the ancestral strain. BA.4/BA.5 breakthrough sera was collected from individuals who had received monovalent mRNA vaccines followed by infection with Omicron sub-lineages BA.4 or BA.5. Samples were examined by anti-nucleoprotein (NP) ELISA to confirm status of prior SARS-CoV-2 infection.

A subset of sera analyzed in this study was collected at the University of Michigan through the Immunity-Associated with SARS-CoV-2 Study (IASO), an ongoing cohort study in Ann Arbor, Michigan that began in 2020[1]. All IASO participants provided written informed consent and serum samples were collected under the protocol approved by the Institutional Review Board of the University of Michigan Medical School.

A subset of vaccinee and breakthrough sera analyzed in this study was collected at Columbia University Irving Medical Center. All subjects provided written informed consent, and all serum collections were performed under protocols reviewed and approved by the Institutional Review Board of Columbia University.

Clinical information for the different study cohorts is summarized in **Table S1** with detailed information on each case provided in **Table S2**.

*Cell lines*

Vero-E6 cells (CRL-1586) and HEK293T cells (CRL-3216) were obtained from the American Type Culture Collection. Cells were maintained in Dulbecco’s Modified Eagle Medium (DMEM) with 10% fetal bovine serum and 1% penicillin-streptomycin in an atmosphere of 5% $CO_2$ at 37 °C.

*SARS-CoV-2 spike plasmids*

Plasmids encoding the spike (S) protein of SARS-CoV-2 variants D614G, BA.1, BA.2, BA.4/BA.5, BA.4.6, BA.2.75, SARS-CoV, GD-Pangolin, GX-Pangolin, and WIV1 were previously constructed[2-7]. BA.2.75.2 spike were constructed with the QuikChange II XL site-directed mutagenesis kit according to the manufacturer's instructions (Agilent). The sequence of each construct was confirmed by Sanger sequencing prior to experimental use.

*Pseudovirus production*

Pseudotyped SARS-CoV-2 variants and other tested sarbecoviruses were generated in the background of vesicular stomatitis virus (VSV). The native VSV glycoprotein (G) was replaced with the S protein from each SARS-CoV-2 variant or other tested sarbecoviruses as previously described[8]. Briefly, HEK293T cells were transfected with plasmids encoding the appropriate S protein using 1 mg/mL of PEI. Transfected HEK293T cells were then cultured at 37 °C with 5% $CO_2$ for 24 hours. Cells were then infected with VSV-G pseudotyped ΔG-luciferase (G*ΔG-luciferase, Kerafast). After a two-hour incubation at 37 °C, infected HEK293T cells were washed three times before being cultured in fresh medium for another 24 hours under the same conditions. Supernatants were subsequently collected, centrifuged to remove precipitates, and aliquoted for storage at -80 °C. Prior to infection of target cells, the viral stock was incubated with 20% I1 hybridoma (anti-VSV-G) supernatant (ATCC; CRL-2700) for 1 h at 37°C to neutralize contaminating VSV-G pseudotyped ΔG-luciferase.

*Pseudovirus neutralization*

Before each neutralization assay, all pseudoviruses were titrated to equilibrate the viral input. Sera were heat-inactivated and all samples run in triplicate in 96-well plates. Sera were four-fold serially diluted in media starting at a 1:100 dilution. Pseudoviruses were added and the virus-sample mixture was incubated at 37 °C for 1 hour. Control wells only containing virus were included on all plates. Vero-E6 cells were then added at a density of $4\times10^4$ cells per well and plates were incubated at 37 °C with 5% $CO_2$ for 10 hours. Cells were then lysed and luciferase activity was measured using the Luciferase Assay System (Promega) and SoftMax Pro v.7.0.2 (Molecular Devices) according to instructions from both manufacturers.

*Quantification and statistical analysis*

The inhibitory dilution retaining 50% neutralization ($ID_{50}$) was obtained for each serum-virus combination using a five-parameter dose-response curve in GraphPad Prism v.9.2. Statistical significance between unpaired groups was evaluated using the two-tailed Mann-Whitney test in GraphPad Prism v.9.2. Levels of significance are denoted as follows: *$p < 0.05$; **$p < 0.01$; and ***$p < 0.001$.

**Acknowledgements**

This study was supported by funding from the NIH SARS-CoV-2 Assessment of Viral Evolution (SAVE) Program as well as the NIH, NIAID contract number 75N93019C00051 (to A.G.). We thank David Manthei, Emily Stoneman, Victoria Blanc, Pamela Bennett-Baker, Savanna Sneeringer, Lauren Warsinske, Theresa Kowalski-Dobson, Alyssa Meyers, Zijin Chu, Hailey Kuiken, Lonnie Barnes, Ashley Eckard, Kathleen Lindsey, Dawson Davis, Aaron Rico, Casey Juntila, Daniel Raymond, Mayurika Patel, and Nivea Vydiswaran from the IASO study team for providing serum samples.

### Author Contributions

L.L. and D.D.H. conceived the study. Q.W. and L.L. performed experiments and analyzed data. Q.W. managed the project. A.B., R.V., C.G. and A.G. collected serum samples. Q.W., A.B., L.L., and D.D.H. analyzed the results and wrote the manuscript. L.L. and D.D.H. directed and supervised the project. All authors reviewed and approved of the manuscript.

### Declaration of Interests

D.D.H. is a co-founder of TaiMed Biologics and RenBio, consultant to WuXi Biologics and Brii Biosciences, and board director for Vicarious Surgical. Aubree Gordon serves on a scientific advisory board for Janssen Pharmaceuticals. Other authors declare no competing interests.

**Table S1. Summary of clinical cohorts**

| Characteristic | 3 shots WT (N=14) | BA.4/BA.5 breakthrough (N=20) | 4 shots WT (N=19) | 3 shots WT + bivalent (N=21) |
|---|---|---|---|---|
| Sex — no. (%) | | | | |
| Female | 6 (42.9%) | 17 (85.0%) | 17 (89.5%) | 16 (76.2%) |
| Male | 8 (57.1%) | 3 (15.0%) | 2 (10.5%) | 5 (23.8%) |
| Mean Age (range) — years | 52.1 (26, 71) | 44.4 (24, 69) | 55.3 (48, 63) | 36.4 (23, 49) |
| Mean days post vaccination or infection (range) | 39.2 (14, 90) | 31.8 (15, 75) | 24.0 (20, 36) | 26.4 (23, 30) |
| First and Second Vaccine Type — no. (%) | | | | |
| Pfizer (BNT162b2) | 12 (85.7%) | 17 (85.0%) | 18 (94.7%) | 17 (81.0%) |
| Moderna (mRNA-1273) | 2 (14.3%) | 3 (15.0%) | 1 (5.3%) | 4 (19.0%) |
| Third Vaccine Type — no. (%) | | | | |
| Pfizer (BNT162b2) | 11 (78.6%) | 15 (75.0%) | 18 (94.7%) | 13 (61.9%) |
| Moderna (mRNA-1273) | 3 (21.4%) | 5 (25.0%) | 1 (5.3%) | 8 (38.1%) |
| Fourth Vaccine Type (monovalent or *bivalent) — no. (%) | | | | |
| Pfizer | - | 10 (50.0%) | 18 (94.7%) | *9 (42.9%) |
| Moderna | - | 0 (0.0%) | 1 (5.3%) | *12 (57.1%) |

### Table S2. Demographics of clinical cohorts

| Sample ID | Vaccine type and infected strain | Days post-vaccination or *infection | Documented COVID-19 | Age | Gender |
|---|---|---|---|---|---|
| 3 shots WT | | | | | |
| Q1 | mRNA-1273/mRNA-1273/mRNA-1273 | 29 | No | 66 | Female |
| Q2 | BNT162b2/BNT162b2/BNT162b2 | 30 | No | 68 | Male |
| Q3 | BNT162b2/BNT162b2/BNT162b2 | 14 | No | 64 | Female |
| Q4 | BNT162b2/BNT162b2/BNT162b2 | 34 | No | 55 | Male |
| Q5 | BNT162b2/BNT162b2/BNT162b2 | 34 | No | 45 | Male |
| Q6 | BNT162b2/BNT162b2/BNT162b2 | 15 | No | 50 | Female |
| Q7 | BNT162b2/BNT162b2/BNT162b2 | 15 | No | 48 | Female |
| Q8 | BNT162b2/BNT162b2/BNT162b2 | 29 | No | 71 | Male |
| Q9 | BNT162b2/BNT162b2/BNT162b2 | 90 | No | 59 | Male |
| Q10 | BNT162b2/BNT162b2/BNT162b2 | 33 | No | 45 | Male |
| Q11 | BNT162b2/BNT162b2/BNT162b2 | 87 | No | 66 | Female |
| Q12 | BNT162b2/BNT162b2/BNT162b2 | 84 | No | 26 | Male |
| Q13 | mRNA-1273/mRNA-1273/mRNA-1273 | 23 | No | 28 | Female |
| Q15 | BNT162b2/BNT162b2/mRNA-1273 | 32 | No | 39 | Male |
| BA.4/BA.5 breakthrough | | | | | |
| Q71 | mRNA-1273/mRNA-1273/BNT162b2/BA.5.2.1 | *29 | Yes | 29 | Female |
| Q77 | BNT162b2/BNT162b2/BNT162b2/BA.5 | *22 | Yes | 61 | Female |
| Q79 | mRNA-1273/mRNA-1273/mRNA-1273/BA.5 | *15 | Yes | 28 | Female |
| Q80 | mRNA-1273/mRNA-1273/mRNA-1273/BA.5 | *21 | Yes | 24 | Female |
| Q81 | BNT162b2/BNT162b2/BNT162b2/BA.5 | *75 | Yes | 35 | Female |
| Q82 | BNT162b2/BNT162b2/mRNA-1273/BA.5 | *63 | Yes | 46 | Female |
| Q83 | BNT162b2/BNT162b2/BNT162b2/BA.5 | *28 | Yes | 55 | Male |
| Q84 | BNT162b2/BNT162b2/BNT162b2/BA.5 | *17 | Yes | 57 | Female |
| UM-85 | BNT162b2/BNT162b2/BNT162b2/BA.5 | *29 | Yes | 44 | Female |
| UM-86 | BNT162b2/BNT162b2/mRNA-1273/BA.5 | *29 | Yes | 36 | Female |
| UM-87 | BNT162b2/BNT162b2/BNT162b2/BNT162b2/BA.5 | *31 | Yes | 54 | Female |
| UM-88 | BNT162b2/BNT162b2/BNT162b2/BNT162b2/BA.5 | *28 | Yes | 69 | Male |
| UM-89 | BNT162b2/BNT162b2/BNT162b2/BNT162b2/BA.5 | *42 | Yes | 44 | Male |
| UM-90 | BNT162b2/BNT162b2/BNT162b2/BNT162b2/BA.5 | *28 | Yes | 41 | Female |
| UM-91 | BNT162b2/BNT162b2/BNT162b2/BNT162b2/BA.5 | *28 | Yes | 44 | Female |
| UM-92 | BNT162b2/BNT162b2/BNT162b2/BNT162b2/BA.5 | *31 | Yes | 29 | Female |
| UM-93 | BNT162b2/BNT162b2/BNT162b2/BNT162b2/BA.5 | *29 | Yes | 48 | Female |
| UM-94 | BNT162b2/BNT162b2/BNT162b2/BNT162b2/BA.5 | *29 | Yes | 49 | Female |
| UM-95 | BNT162b2/BNT162b2/mRNA-1273/BNT162b2/BA.5 | *28 | Yes | 37 | Female |
| UM-96 | BNT162b2/BNT162b2/BNT162b2/BNT162b2/BA.5 | *33 | Yes | 58 | Female |
| 4 shots WT | | | | | |
| UM-65 | BNT162b2/BNT162b2/BNT162b2/BNT162b2 | 24 | No | 52 | Female |
| UM-66 | BNT162b2/BNT162b2/BNT162b2/BNT162b2 | 20 | No | 57 | Female |
| UM-67 | BNT162b2/BNT162b2/BNT162b2/BNT162b2 | 20 | No | 61 | Female |
| UM-68 | mRNA-1273/mRNA-1273/mRNA-1273/mRNA-1273 | 22 | No | 48 | Female |
| UM-69 | BNT162b2/BNT162b2/BNT162b2/BNT162b2 | 23 | No | 50 | Female |
| UM-70 | BNT162b2/BNT162b2/BNT162b2/BNT162b2 | 22 | No | 50 | Female |
| UM-71 | BNT162b2/BNT162b2/BNT162b2/BNT162b2 | 20 | No | 58 | Female |
| UM-72 | BNT162b2/BNT162b2/BNT162b2/BNT162b2 | 26 | No | 56 | Female |
| UM-73 | BNT162b2/BNT162b2/BNT162b2/BNT162b2 | 29 | No | 63 | Female |
| UM-74 | BNT162b2/BNT162b2/BNT162b2/BNT162b2 | 25 | No | 58 | Female |
| UM-75 | BNT162b2/BNT162b2/BNT162b2/BNT162b2 | 21 | No | 62 | Male |
| UM-76 | BNT162b2/BNT162b2/BNT162b2/BNT162b2 | 26 | No | 54 | Female |
| UM-77 | BNT162b2/BNT162b2/BNT162b2/BNT162b2 | 23 | No | 53 | Male |
| UM-78 | BNT162b2/BNT162b2/BNT162b2/BNT162b2 | 21 | No | 55 | Female |
| UM-79 | BNT162b2/BNT162b2/BNT162b2/BNT162b2 | 23 | No | 59 | Female |
| UM-80 | BNT162b2/BNT162b2/BNT162b2/BNT162b2 | 21 | No | 49 | Female |
| UM-81 | BNT162b2/BNT162b2/BNT162b2/BNT162b2 | 27 | No | 57 | Female |
| UM-82 | BNT162b2/BNT162b2/BNT162b2/BNT162b2 | 27 | No | 55 | Female |
| Q97 | BNT162b2/BNT162b2/BNT162b2/BNT162b2 | 36 | No | 53 | Female |
| 3 shots WT + bivalent | | | | | |
| UM-36 | BNT162b2/BNT162b2/BNT162b2/Moderna Bivalent | 24 | No | 38 | Female |

| Sample ID | Vaccine type and infected strain | Days post-vaccination or *infection | Documented COVID-19 | Age | Gender |
|---|---|---|---|---|---|
| UM-37 | BNT162b2/BNT162b2/BNT162b2/Moderna Bivalent | 27 | No | 42 | Female |
| UM-39 | mRNA-1273//mRNA-1273/mRNA-1273/Moderna Bivalent | 24 | No | 36 | Male |
| UM-40 | BNT162b2/BNT162b2/BNT162b2/Pfizer Bivalent | 25 | No | 37 | Female |
| UM-41 | BNT162b2/BNT162b2/BNT162b2/Pfizer Bivalent | 24 | No | 36 | Male |
| UM-43 | BNT162b2/BNT162b2/BNT162b2/Pfizer Bivalent | 25 | No | 49 | Female |
| UM-44 | BNT162b2/BNT162b2/BNT162b2/Moderna Bivalent | 25 | No | 37 | Female |
| UM-47 | BNT162b2/BNT162b2/BNT162b2/Pfizer Bivalent | 26 | No | 45 | Male |
| UM-48 | BNT162b2/BNT162b2/mRNA-1273/Moderna Bivalent | 26 | No | 43 | Female |
| UM-51 | mRNA-1273/mRNA-1273/mRNA-1273/Moderna Bivalent | 29 | No | 32 | Female |
| UM-52 | BNT162b2/BNT162b2/BNT162b2/Pfizer Bivalent | 23 | No | 43 | Female |
| UM-53 | BNT162b2/BNT162b2/BNT162b2/Pfizer Bivalent | 26 | No | 43 | Female |
| UM-54 | BNT162b2/BNT162b2/mRNA-1273/Moderna Bivalent | 29 | No | 38 | Female |
| UM-55 | BNT162b2/BNT162b2/BNT162b2/Moderna Bivalent | 28 | No | 38 | Female |
| UM-56 | BNT162b2/BNT162b2/mRNA-1273/Moderna Bivalent | 27 | No | 36 | Female |
| UM-60 | BNT162b2/BNT162b2/BNT162b2/Moderna Bivalent | 30 | No | 24 | Female |
| Q101 | mRNA-1273/mRNA-1273/mRNA-1273/Moderna Bivalent | 30 | No | 32 | Female |
| Q102 | BNT162b2/BNT162b2/mRNA-1273/Moderna Bivalent | 23 | No | 39 | Male |
| Q103 | BNT162b2/BNT162b2/BNT162b2/Pfizer Bivalent | 30 | No | 26 | Female |
| Q104 | mRNA-1273/mRNA-1273/mRNA-1273/Pfizer Bivalent | 30 | No | 27 | Female |
| Q105 | BNT162b2/BNT162b2/BNT162b2/Pfizer Bivalent | 23 | No | 23 | Male |
